# CRISPR-LRS for mapping transgenes in the mouse genome

**DOI:** 10.1101/2022.01.05.475144

**Authors:** W. Bart Bryant, Allison Yang, Susan Griffin, Wei Zhang, Xiaochun Long, Joseph M. Miano

## Abstract

Microinjected transgenes, including bacterial artificial chromosomes (BACs), insert randomly in the mouse genome. Traditional methods of mapping a transgene are challenging, thus complicating breeding strategies and the accurate interpretation of phenotypes, particularly when a transgene disrupts critical coding or noncoding sequences. Here, we introduce CRISPR-Cas9 long-read sequencing (CRISPR-LRS) to ascertain transgene integration locus and estimated copy number. This method revealed integration loci for both a BAC and *Cre*-driver line, and estimated the copy numbers for two other BAC mouse lines. CRISPR-LRS offers an easy approach to establish robust breeding practices and accurate phenotyping of most any transgenic mouse line.

---

Despite the routine use of transgenic mice, greater than 90% of transgenic alleles in the Mouse Genome Database have yet to be mapped (1, 2) even though it is broadly accepted that random integration of a transgene can disrupt coding exons and functional noncoding sequences (eg, enhancer, long noncoding RNA),thus complicating data interpretation. Fluorescence in situ hybridization (FISH) can map a transgene to a chromosomal band, but lacks nucleotide resolution (3). Hybridization, targeted locus amplification (TLA), and linear amplification-mediated PCR (LAM-PCR), rely on short-read sequencing and cannot resolve complex genome inversions/deletions (4–7). Inverse PCR has high failure rates due to concatemerization of most transgenes (8). Recently, whole genome sequencing (WGS) with long-read sequencing platforms, PacBio or Oxford Nanopore Technologies (ONT), have been used to map transgenes (2, 9). However, even with ~7.5Gb of sequencing data for ~3x coverage of the mouse genome, researchers cannot be assured to find reads that identify the breakpoint of the transgene.

We propose a more targeted approach to map a transgene by combining (i) the RNA programmable CRISPR-Cas9 system (10) and (ii) long-read length coverage of the ONT sequencing platform (11, 12). Previous studies have combined CRISPR-Cas9 with LRS to enrich for genomic elements (13–20); however, no studies until now have combined CRISPR and LRS to map transgenes in an animal model. CRISPR-LRS can be designed (i) to target a genomic section (Targeted-CRISPR-LRS) or (ii) to enrich genomic sections (Enrichment-CRISPR-LRS). In this study, CRISPR-LRS successfully mapped a single-copy BAC and a multi-copy *Cre* in the mouse genome. CRISPR-LRS represents a facile tool for mapping transgenes in experimental animal models, thus informing investigators as to best breeding practices and potential genetic confounders.

## Results/Discussion

For the four transgenic mouse lines in this study, seven CRISPR-LRS libraries (Targeted or Enrichment) were sequenced with the minION platform for a total of ~1.8Gb at ~400,000 reads (see Supp. Table 1 and 2). Reads mapped to their corresponding reference sequence with a range of 0.02% - 0.52% (Supp. Table 2) and will be referred to as informative long-reads below.

### CRISPR-LRS mapping of BAC mouse lines

The mouse line carrying the RP11-744N12 BAC was generated to study human-specific long noncoding RNA, *SENCR* (21). To map where this 217 kb BAC integrated into the mouse genome, two Targeted-CRISPR-LRS libraries were made 5 kb or 3 kb from the 5’ and 3’ ends of each BAC sequence, respectively (Figure 1A, Supp. Fig 1i, and Supp. Table 1). Both libraries yielded 0.03% of reads (0.9 kb – 25 kb) that mapped to the reference sequence (Figure 1Ai and Supp. Table 2). Notably, manual inspection of informative long-reads revealed ~7 kb of pBACe3.6 vector sequence accompanying transgene and mouse chromosome 15 sequences (Figure 1Ai). Further, informative long-reads over 6 kb for 5’ end and 10 kb for 3’ end, found RP11-744N12 integrated within the first intron of *Egflam* (Chr15:7,344,678; GRCm38/mm10) (Figure 1A). CRISPR-LRS also mapped mouse BAC lines, CTD-2518N7 and RP11-997L11, and found informative long-reads with the BAC-cloning vector flanked by 5’ and 3’ human sequence, indicating integration of at least two copies of each transgenic line (Supp. Fig 1ii and Supp. Fig 2).

**Figure 1.**
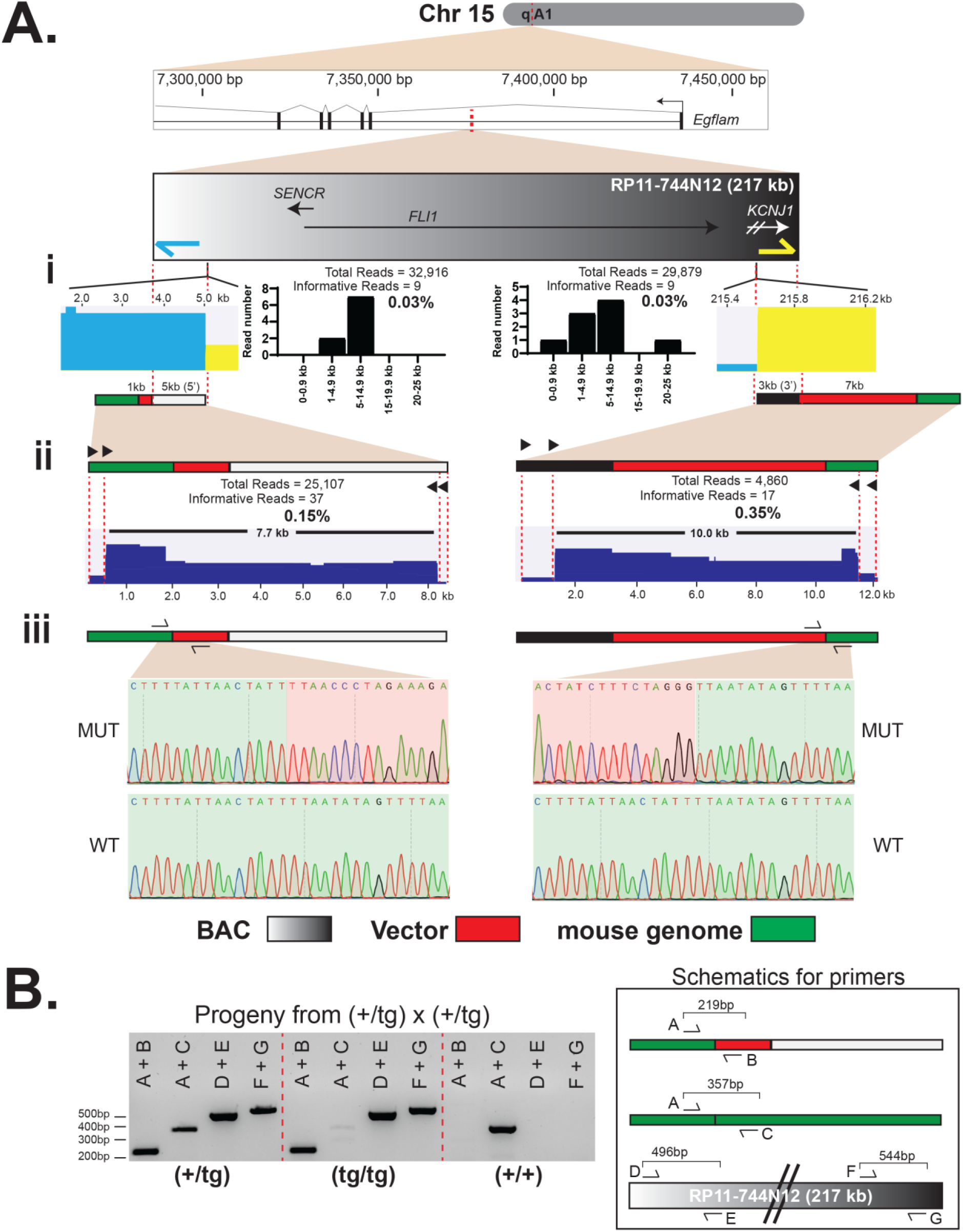
CRISPR-LRS mapping of RP11-744N12 BAC to chromosome 15 in mouse genome. (A) Dashed red line represents integration locus of RP11-744N12 BAC within first intron of *Egflam* on chromosome 15, section qA1. Blue and yellow arrows represent the 5’ and 3’ end crRNAs, respectively, used to make Targeted-CRISPR-LRS libraries (see Supp. Fig1i flow chart). (i) Histograms for informative reads from Targeted-CRISPR-LRS libraries at either 5’ or 3’ end of the RP11-744N12 BAC. (ii) Black triangles represent flanking tandem crRNAs used to make Enrichment-CRISPR-LRS libraries to interrogate 5’ and 3’ ends of the RP11-744N12 integration locus on chromosome 15, section qA1. (iii) Sanger sequencing verification of CRISPR-LRS mapped RP11-744N12 integration locus for 5’ and 3’ ends of the RP11-744N12 integration locus. (B) PCR genotyping of mice progeny from (+/tg) x (+/tg) cross with primer schematics. Primer pair A+B interrogated the breakpoint junction of RP11-744N12 BAC and chromosome 15, section qA1. Primer pair A+C interrogated wild type integration locus. To check for presence of RP11-744N12 BAC transgene, internal primer pairs D+E and F+G targeted 5’ and 3’ regions of the RP11-744N12 BAC sequence, respectively.

While Targeted-CRISPR-LRS mapped the RP11-744N12 BAC integration locus, only a few long-reads covered the integration loci. Enrichment-CRISPR-LRS queried 7.7 kb and 10.0 kb for the 5’ and 3’ terminal ends, respectively, with crRNAs enriching for (i) mouse chromosome 15, (ii) pBACe3.6 cloning vector, and (iii) RP11-744N12 sequences (Figure 1Aii and Supp. Fig 1iv). With 5- and 12-fold enrichment of informative reads at the 5’- and 3’-terminal ends, respectively (compare Figure 1Ai to 1Aii), Enrichment-CRISPR-LRS validated the Targeted-CRISPR-LRS mapping data (Figure 1Aii). Further, Sanger sequencing confirmed the breakpoint of RP11-744N12 BAC and chromosome 15, within the first intron of *Egflam* (Figure 1Aiii and Supp. Table 1

Genotyping pups from a (+/tg) x (+/tg) cross with primers spanning (i) transgene and chromosome 15 breakpoint and (ii) wild type chromosome 15 loci (Figure 1B), revealed near Mendelian ratio: (+/tg), 17/41 at 41%; (+/+), 9/41 at 22%; (tg/tg), 15/41 at 37%. Notably, the BAC transgene exists as a single copy with no loss of mouse genomic sequence at the site of integration and healthy, homozygous transgenic mice indicate the absence of overt pathology.

While CRISPR-LRS mapped RP11-744N12 as a single copy, we wanted to check for the possibility of additional integration sites as one study reported multiple integration loci for ~10% of their EGFP-reporter lines by FISH (3). To verify a single integration locus, heterozygous (+/tg) pups were crossed and progeny genotypes were assessed with BAC-specific primers, (Figure 1B). As expected, BAC-specific amplicons were present in (+/tg) and (tg/tg) pups (Figure 1B). However, if a wild type (+/+) pup from the heterozygous cross exhibited BAC-specific amplicons, then this result would indicate that CRISPR-LRS missed additional transgene integration sites. Notably, wild type (+/+) pups from the (+/tg) x (+/tg) did not exhibit BAC-specific amplicon products (Figure 1B), demonstrating CRISPR-LRS did successfully map the RP11-744N12 BAC as a single copy transgenic mouse line.

Fortuitously, as all BAC transgenes were flanked with BAC-cloning vector sequence (Figure 1 and Supp. Fig 2), copy number quantification was possible. qPCR, routine for copy number variation analysis (1, 2, 5, 9), targeted the *chloramphenicol resistance gene*, a common gene within BAC-vectors. With a single integration (Figure 1), (+/tg) and (tg/tg) RP11744N12 pups were queried by qPCR finding one and two BAC transgene copies, respectively (Supp. Fig 3A). As the other BAC-lines exhibited tandem integrations, only (+/tg) pups were queried, with qPCR showing ~2-3 copies for both lines. Collectively, qPCR was consistent with the CRISPR-LRS mapping data (Figure 1, Supp. Fig1 i and ii, Supp. Fig 2, and Supp. Fig 3A).

### CRISPR-LRS mapping of *Cre*-driver mouse line

Most small transgenes lack a mapped integration locus and can possess dozens of copies of the transgene, as well as complex rearrangements of the host genome (1, 2, 22). We next applied CRISPR-LRS to map *Sm22-Cre*, a mouse line used to excise floxed DNA sequences in early embryonic heart and smooth muscle cell-containing tissues (23). To map the BAC lines, CRISPR-LRS targeted at least 2 kb from terminal ends of the transgene (Figure 1, Supp. Fig 1i-ii, and Supp. Fig 2). However, because *Cre* is significantly smaller than a BAC, a different Targeted-CRISPR-LRS approach was used. Specifically, two independent libraries targeting 0.8 kb or 0.5 kb from the 5’ and 3’ end, respectively, were run on one flow cell (Figure 2Ai, Supp. Fig 1iii, and Supp. Table 1). At 0.52%, the *Sm22-Cre* libraries contained more informative long-reads over the BAC libraries (Figure 1, Figure 2Ai, Supp. Fig 2, and Supp. Table 2). Manual interrogation and alignment of >6 kb informative long-reads elucidated a mini-tiled *Cre* transgene integration map consisting of multiple copies of *Cre* and genomic inversions (Figure 2Ai). Three informative long-reads revealed a breakpoint between one of the *Cre* transgenes and 91,527,881bp on chromosome 14 (GRCm38/mm10) (Figure 2Ai, dashed black line box). Sanger sequencing verified the *Cre* and host chromosome breakpoint (Figure 2Aii and Supp. Table 1).

**Figure 2.**
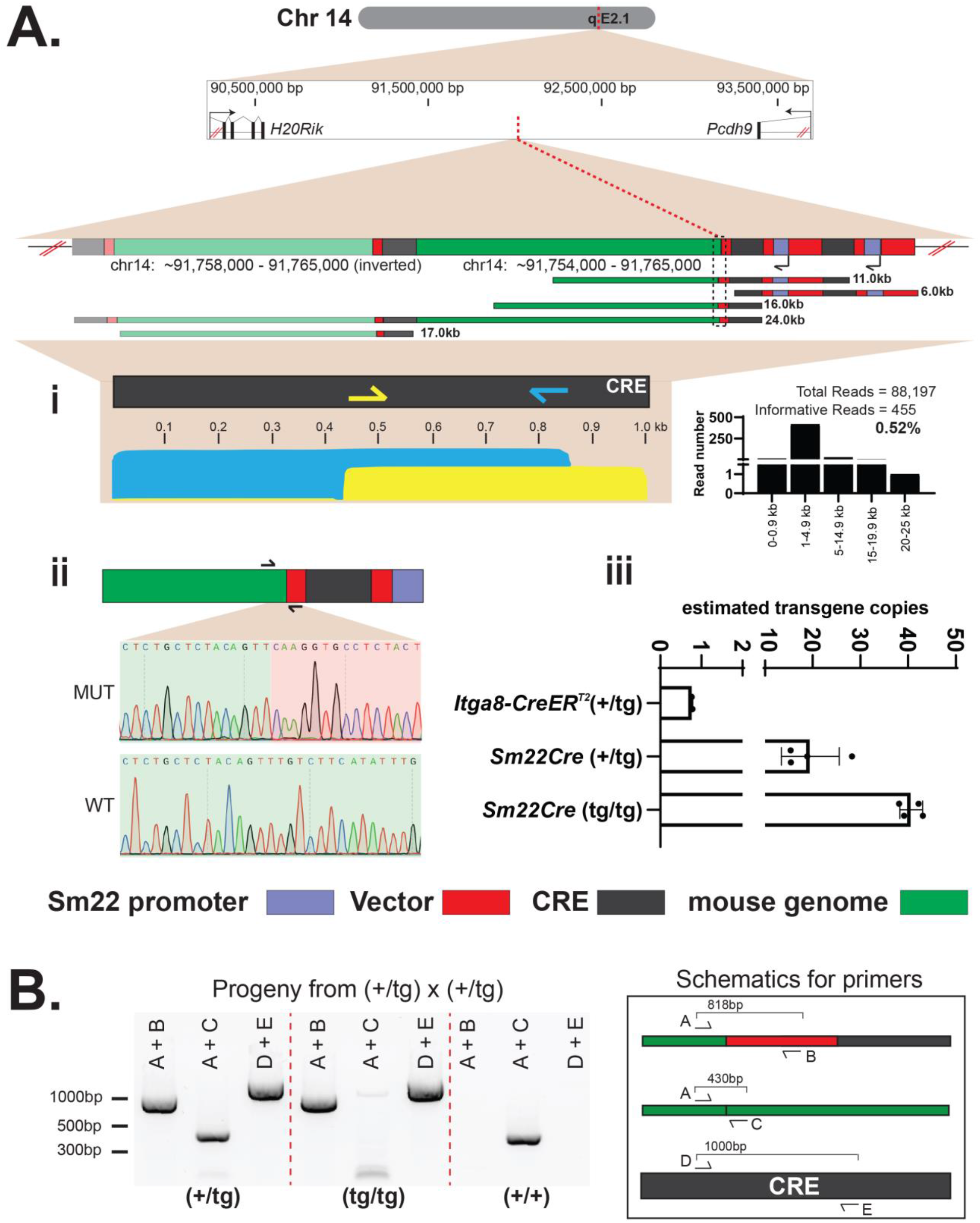
CRISPR-LRS mapping of *Sm22Cre*-driver to chromosome 14 in mouse genome. (A) Dashed red line represents integration locus of *Sm22Cre*-driver within chromosome 14, section qE2.1. Informative long reads were compiled to build an integration locus map. Dashed black line box represents *Sm22Cre*-driver integration locus. Blue and yellow arrows represent internal overlapping 5’ and 3’ end crRNAs used to make a Targeted-CRISPR-LRS library (see Supp. Fig 1iii flow chart). (i) Histogram for informative reads from the Targeted-CRISPR-LRS library. (ii) Sanger sequencing verification of CRISPR-LRS mapped *Sm22Cre*-driver integration locus, represented by dashed black line box highlighted in (A). (iii) qPCR determination of transgene copy number with (+/tg) and (tg/tg) pups with approximately 20 and 40 copies, respectively. Data normalized to internal control locus and calibrator (*Itga8-CreER^T2^*, see Methods for details). *N* = 2 for calibrator (*Itga8-CreER^T2^*), *N* = 4 for *Sm22Cre* (+/tg) and (tg/tg). Values graphed as mean ± SD. (B) PCR genotyping of mice progeny from (+/tg) x (+/tg) cross with primer schematics. Primer pair A+B interrogated the breakpoint junction of *Sm22Cre*-driver and chromosome 14, section qE2.1, represented by dashed black line box highlighted in (A). Primer pair A+C interrogated wild type integration locus. To check for presence of transgene, internal primer pair D+E targeted the *Cre* transgene.

Genotyping pups from a (+/tg) x (+/tg) cross with primers spanning (i) the breakpoint of *Cre* and chromosome 14 and (ii) wild type chromosome 14 loci (Figure 2B), revealed near Mendelian ratio: (+/tg), 20/41 at 49%; (+/+), 8/41 at 19%; (tg/tg), 13/41 at 32%.

To check for additional integration loci for the *Cre*-driver, heterozygous pups were crossed and progeny genotypes assessed with *Cre*-specific primers, D and E (Figure 2B). Both (+/tg) and (tg/tg) pups yielded amplicon products for the internal *Cre* primers, as expected (Figure 2B). Notably, wild type pups from the (+/tg) x (+/tg) cross did not yield *Cre*-specific amplicon products (Figure 2B), demonstrating CRISPR-LRS successfully mapped one integration locus for the *Sm22Cre* mouse line.

As small transgenes typically integrate as concatemers (1, 2), our mini-tiled integration map for *Sm22Cre* could not firmly establish copy number (Figure 2A). To address this limitation, qPCR determined the copy number for the line with ~20 and ~40 copies of *Cre* for (+/tg) and (tg/tg) pups, respectively (Figure 2Aiii and Supp. Figure 3B). As the *Sm22Cre* mouse has ~20 copies of *Cre* per allele, there were more Cas9-ribonucleoprotein (RNP) targets to cleave compared to the BAC lines, explaining how the *Sm22Cre*-CRISPR-LRS libraries contained more informative mapped long-reads over the BAC-CRISPR-LRS libraries (Figure 1, Figure 2, Supp. Fig 2, Supp. Fig 3, and Supp. Table 2).

## Conclusions

There are several benefits of mapping transgenes in animal models such as mice. First, mapping allows investigators to accelerate the generation of desired outcomes in a complex breeding scheme, (e.g., floxed alleles with *Cre*-driver lines). Second, it enables quality design of genotyping assays for colony maintenance and zygosity determination. Lastly, and arguably most important, it alerts one to confounding genetics caused by insertion of the transgene in a coding/noncoding sequence or regulatory element (1, 2, 5, 9, 22).

Of the available strategies to map transgenes, WGS (2, 9) and TLA (1, 5) have the most traction. However with WGS, most of the sequence data is uninformative and sequence depth is substantial with ~2-5x coverage of the host genome (2, 9). Here, with only ~1.8Gb of cumulative sequence data, CRISPR-LRS mapped four mouse lines with two lines yielding PCR validated chromosome coordinates. TLA can be technically challenging for most labs as (i) it has numerous steps; crosslinking, fragmentation, re-ligation, and amplification, and (ii) analysis can require extensive computational expertise due to the nature of these fragmented sequence libraries (1, 5). CRISPR-LRS (i) has less steps; Cas9 cleavage and adaptor ligation, and (ii) only requires mapping long-reads with an open source tool, minimap2 (24). We envision CRISPR-LRS as the ‘go-to’ method of mapping transgenes in any organism with a reference genome.

## Methods

### Transgenic mice

The *SENCR* BAC and *Sm22Cre* mouse lines were reported previously (21), (23). The human CTD-2518N7 and RP11-997L11 BAC transgenic lines were generated by Cyagen (www.cyagen.com) using strain C57BL/6J. All mice were maintained on strain C57BL/6J through repeated back-crossing, refreshing the breeders every 5 generations to mitigate genetic drift. Mouse experiments were approved by Medical College of Georgia at Augusta University Institutional Animal Care and Use Committee (approval numbers 2019-1000 and 2019-0999).

### Long-read library preparation

Genomic DNA (gDNA) was isolated from mouse liver tissue using Qiagen DNeasy Blood & Tissue Kit (cat#69504) following manufacturer’s instructions (www.qiagen.com). To limit shearing of gDNA, wide bore pipette tips were used. Libraries (5μg gDNA) were prepared for four different transgenic mouse lines following manufacturer’s instructions for Cas9 sequencing kit (SQK-CS9109) using the long fragment buffer option during library prep clean-up for seven total CRISPR-LRS libraries (www.nanoporetech.com). crRNAs were designed using CHOPCHOP (25) with default parameters (Supp. Table I) (https://chopchop.cbu.uib.no). Following suggestions from ONT, all crRNAs, tracrRNA, and HiFi Cas9 were ordered from IDT (www.idt.dna.com). To ensure adequate read length needed to accurately map transgene integration loci, crRNAs were designed to target within 5 kb of the terminal 5’ and 3’ ends of the BAC sequence. Further, a Targeted- or an Enrichment-CRISPR-LRS approach was performed for each mouse line (see Supp. Fig. 1 for flow chart). For Targeted-CRISPR-LRS, crRNAs were designed at the 5’ and 3’ ends of the transgene where RNPs were loaded with either one or multiple crRNAs (Supp. Figure 1i - iii and Supp. Table 1). For Enrichment-CRISPR-LRS, tandem crRNAs were designed up- and down-stream of the genomic region of interest (ROI) and loaded onto one RNP (14, 20) (Supp. Figure 1iv and Supp. Table 1). For both approaches, pre-existing DNA ends were dephosphorylated before Cas9-cutting, which yielded preferential ligation of nanopore adaptors to fresh Cas9 cleavage sites as a means to target specific genomic ROIs. For the RP11-744N12 BAC mouse line, two crRNAs, one targeting the 5’ and one targeting the 3’ end of the BAC sequence, were loaded onto two separate RNPs for two independent libraries (Supp. Fig. 1i). Since an integration locus for the RP11-744N12 BAC mouse line was determined, Enrichment-CRISPR-LRS was further performed following manufacturer’s instructions (SQK-CS9109) (www.nanoporetech.com). Both 5’ and 3’ integration loci were probed with four crRNAs loaded onto one RNP (Supp. Fig. 1iv). For the remaining BAC mouse lines, CTD-2518N7 and RP11-997L11, two crRNAs were loaded onto one RNP for one Cas9 library run (Supp. Fig. 1ii). For the *Cre*-driver mouse line, overlapping crRNAs targeting *Cre* were designed at least 0.5 kb from 5’ or 3’ end with two crRNAs loaded onto two separate RNPs and the two independent libraries combined on one flow cell (Supp. Fig. 1iii).

### Nanopore sequencing and data analysis

Cas9 targeted long-read libraries were run on R9.4.1 flow cells on a minION Mk 1B following manufacturer’s instructions for Cas9 sequencing kit (SQK-CS9109) (www.nanoporetech.com). Reads were converted from fast5 to fastq with guppy (v4.2.2) on MinKNOW (v20.10.3) MinKNOW Core (v4.1.2) with fast base-calling option for the base-call model and minimum Q-score of 7 option for read filtering. To analyze LRS results, guppy base-called fastq files were imported into Qiagen CLC Genomics Workbench (www.qiagen.com).Reference sequences specific to each transgenic mouse line were obtained from NCBI nucleotide database (www.ncbi.nlm.nih.gov/nucleotide/)and alignments generated using the Long-Read Support (beta) plugin available in Qiagen CLC Genomics Workbench (www.digitalinsights.qiagen.com), which utilizes components of open-source tool minimap2 (24). Default parameters for the long-read alignment plugin were used. Aligned informative long-reads were extracted and manually queried against NCBI nr/nt and refseq genome databases (https://blast.ncbi.nlm.nih.gov/Blast.cgi) and UCSC genome browser with BLAT tool (https://genome.ucsc.edu). Graphical output obtained from CLC Genomics Workbench and GraphPad (www.graphpad.com) were amended with Adobe Illustrator (www.adobe.com) (Adobe Systems, San Jose, CA, USA) for illustration purposes.

### Genotyping and transgene copy number analysis

Small ear biopsies were taken before weaning and gDNA was extracted using Qiagen DNeasy Blood & Tissue Kit (cat#69504) following manufacturer’s instructions (www.qiagen.com). Progeny of (+/tg) x (+/tg) heterozygous crosses were assessed for CRISPR-LRS mapped integration loci of the transgene. For genotyping, PCR conditions were the following: step 1, 95 °C for 3 min; step 2, 95 °C for 30 sec, 58 °C for 30 sec, and 72 °C for 1 min for 35 cycles; step 3, 72 °C for 10 min. Sequences for all genotyping amplicons were confirmed by Sanger sequencing (Supp. Table I). For transgene copy number determination, gDNA was diluted to 50ng for input. For the BAC mouse lines, two primer sets to the *chloramphenicol resistance gene* served as proxy for the BAC transgene, where values were normalized to an internal control locus (Supp. Table 1) (1, 2, 9). For the *Cre*-driver mouse line, two primer sets to *Cre* were normalized to same internal control locus used for the BAC mouse lines. The *Itga8-CreER^T2^* mouse, known to have one copy of *Cre* (manuscript in preparation), served as a calibrator for one copy of *Cre*. Real time quantitative PCR conditions were the following: step1, 95 °C for 3 min; step 2, 95 °C for 30sec, 60 °C for 30 sec, and 72 °C for 30 sec for 40 cycles.

## Data availability

Data generated by ONT LRS have been submitted to NCBI SRA database (www.ncbi.nlm.nih.gov/sra) under BioProject number PRJNA759232 and will be publically available after manuscript acceptance.

## Declarations

### Ethics approval and consent to participate

Not applicable

### Consent for publication

Not applicable

### Availability of data and materials

Nanopore long read sequencing data are available at NCBI Sequence Read Archive (SRA) under accession number PRJNA759232.

### Competing interests

The authors declare no competing interests

### Funding

Work was supported by grants HL138987 and HL147476 to J.M.M. and HL122686 and HL139794 to X.L.

### Author contributions

W.B.B. and J.M.M designed the study.

A.Y., S.G. and WZ maintained mouse colonies.

W.B.B. performed the experiments.

W.B.B. and J.M.M. analyzed and interpreted data

X.L. provided liver tissue.

W.B.B. and J.M.M. wrote the paper.

All authors read and approved the final manuscript.

## Acknowledgements

We wish to thank Akelia Wauchope-Odumbo and Carl Woodham at Oxford Nanopore Technologies (ONT) for their combined help with use of long-read nanopore sequencing.

---

**Supplemental Figure 1.**
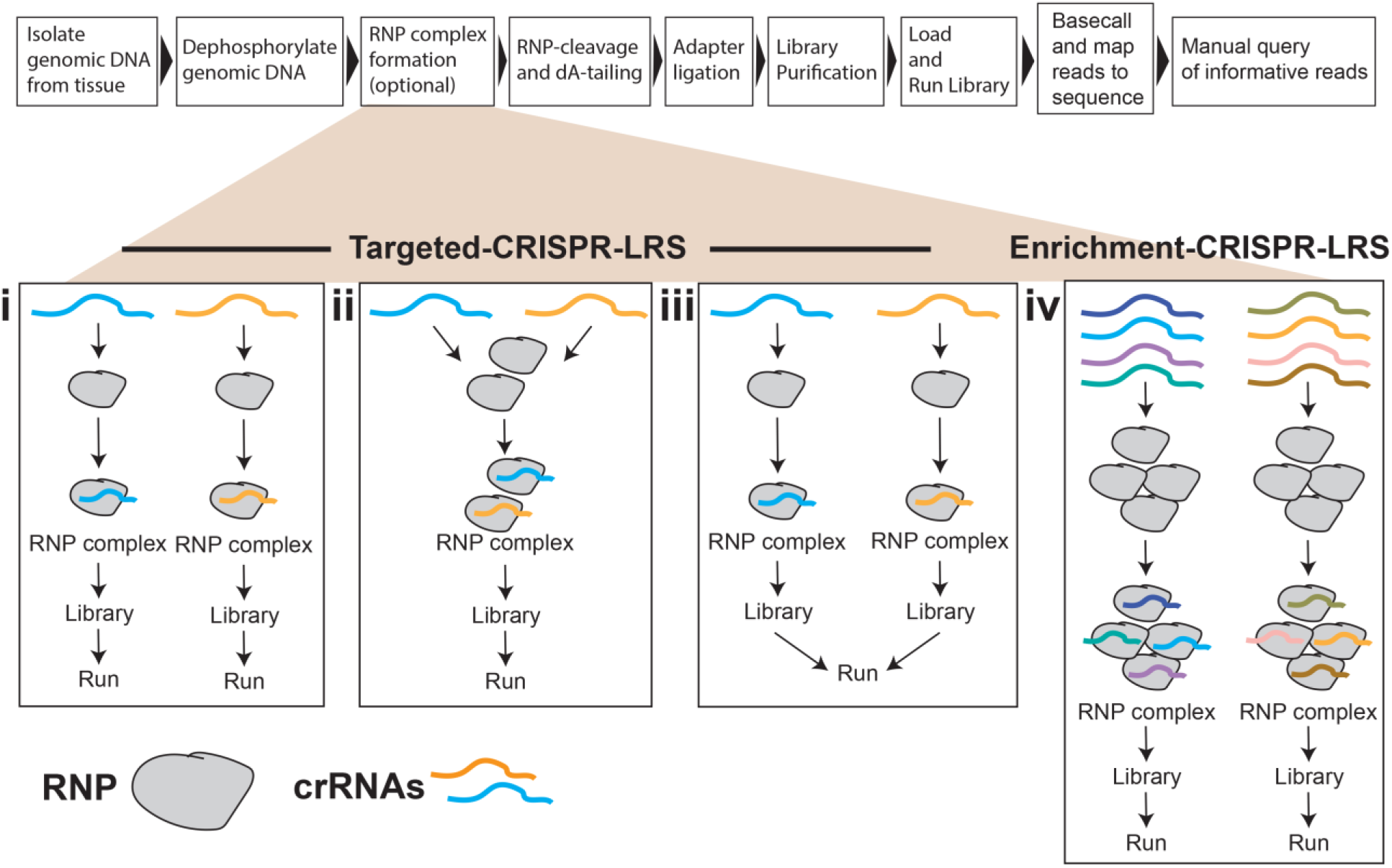
CRISPR-LRS flow chart. Overview of library preparation for Targeted- (i-iii) and Enrichment- (iv) CRISPR-LRS libraries.

**Supplemental Figure 2.**
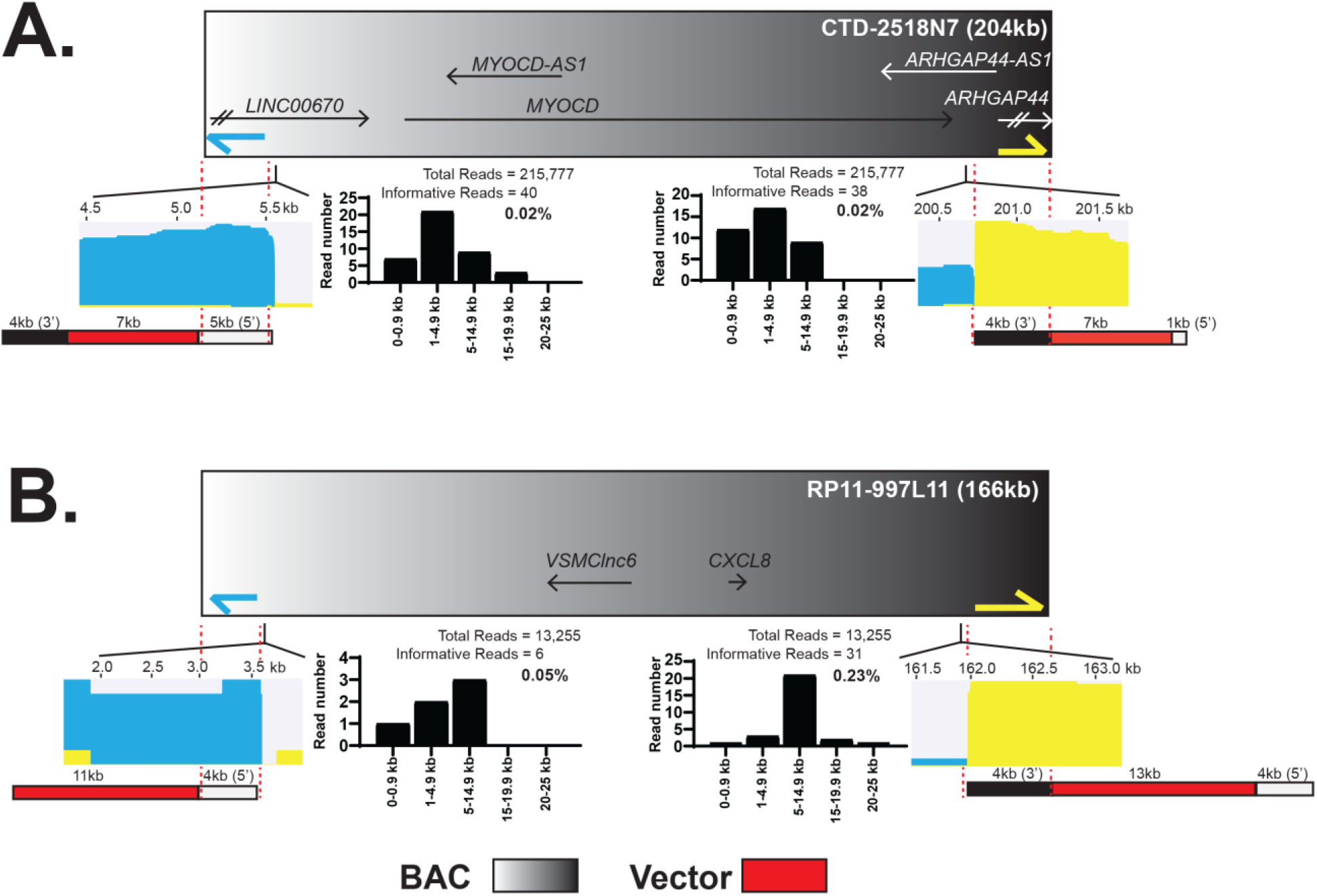
CRISPR-LRS determination of tandem BAC integration for two BAC mouse lines. Overview of Targeted-CRISPR-LRS sequencing illustrating tandem integration of BAC sequence for (A) CTD-2518N7 and (B) RP11-997L11 BAC mouse lines (See Supp. Fig1ii flow chart).

**Supplemental Figure 3.**
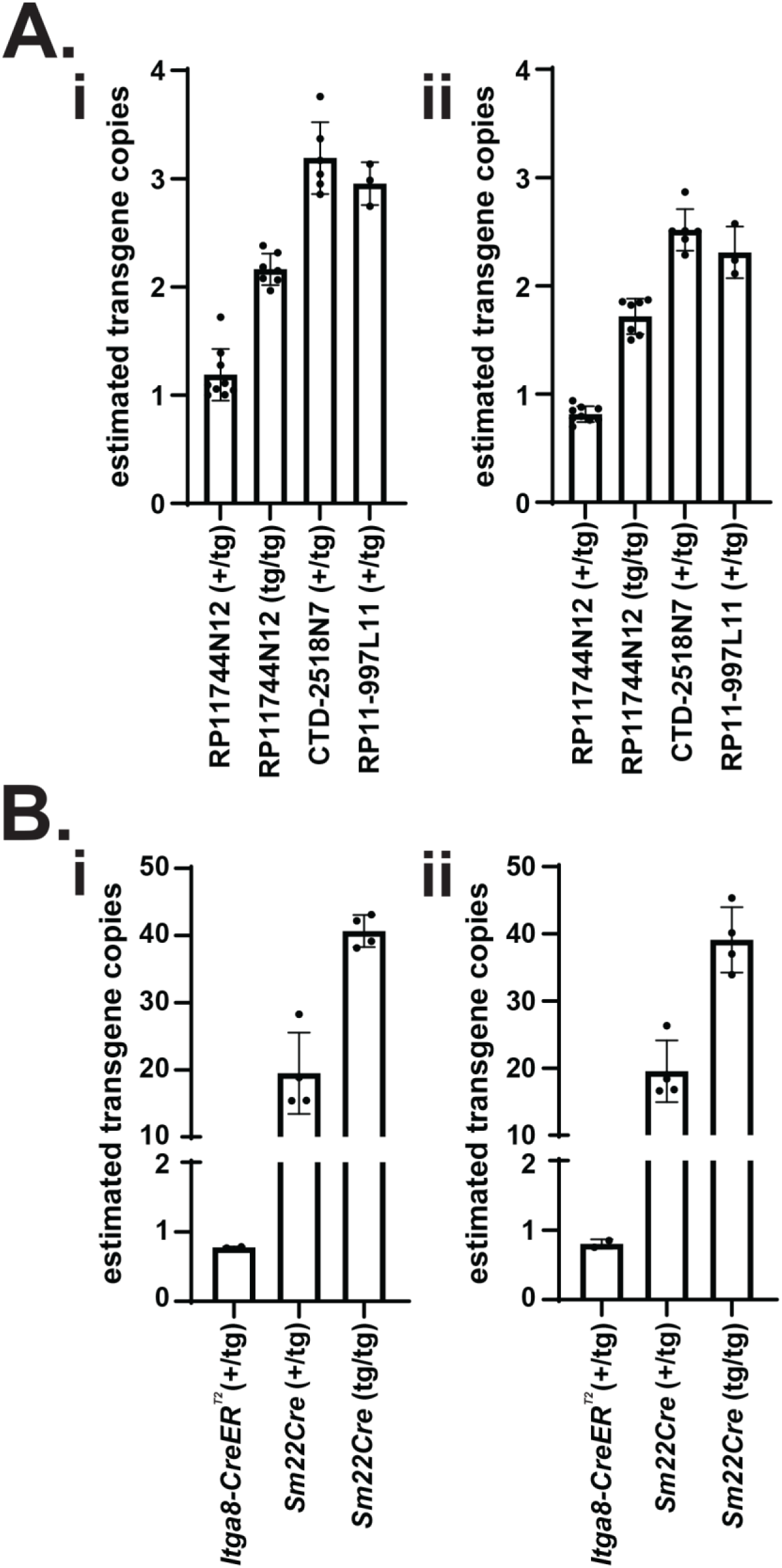
Relative transgene copy number for BAC and *Cre*-driver mouse lines. qPCR to determine transgene copy number, normalized to internal control locus. (A) Two primer sets (i and ii) targeting *chloramphenicol resistance gene*, common gene found in BAC cloning vectors. For RP11-744N12 mouse line, (+/tg) and (tg/tg) pups demonstrated one and two copies, respectively. For both CTD-2518N7 and RP11-997L11 mouse lines, (+/tg) pups demonstrated approximately 3 copies. (B) Two primer sets (i and ii) targeting *Cre* sequence. *Itga8-CreER^T2^* mouse served as calibrator for one copy of *Cre*, serving as (+/tg) control. For *Sm22Cre*-driver mouse line, (+/tg) and (tg/tg) pups demonstrated approximately 20 and 40 copies, respectively. *n* = 9 for RP11-744N12 (+/tg), *n* = 7 for RP11-744N12 (tg/tg), *n* = 6 for CTD-2518N7 (+/tg), *n* = 3 for RP11-997L11 (+/tg), *n* = 2 for *Itga8-CreER^T2^*, *n* = 4 for both *Sm22Cre* (+/tg) and (tg/tg). Values graphed as mean ± SD.

**Supplemental Table 1. crRNAs and primers used for CRISPR-LRS**

**Supplemental Table 2. Nanopore long-read library metrics for CRISPR-LRS**

## Notes

### Competing Interest Statement

The authors have declared no competing interest.

